# The discovery of potential natural products for targeting SARS-CoV-2 spike protein by virtual screening

**DOI:** 10.1101/2020.06.25.170639

**Authors:** Guan-Yu Chen, Tsung-You Yao, Azaj Ahmed, Yi-Cheng Pan, Juan-Cheng Yang, Yang-Chang Wu

## Abstract

Severe acute respiratory syndrome coronavirus 2 (SARS-CoV-2) enters into the cells through its spike proteins binding to human angiotensin-converting enzyme 2 (ACE2) protein and causes virus infection in host cells. Until now, there are no available antiviral drugs have been reported that can effectively block virus infection. The study aimed to discover the potential compounds to prevent viral spike proteins to bind to the human ACE2 proteins from Taiwan Database of Extracts and Compounds (TDEC) by structure-based virtual screening. In this study, to rapidly discover potential inhibitors against SARS-CoV-2 spike proteins, the molecular docking calculation was performed by AutoDock Vina program. Herein, we found that 39 potential compounds may have good binding affinities and can respectively bind to the viral receptor-binding domain (RBD) of spike protein in the prefusion conformation and spike-ACE2 complex protein *in silico*. Among those compounds, especially natural products thioflexibilolide A and candidine that were respectively isolated from the soft corals *Sinularia flexibilis* and *Phaius mishmensis* may have better binding affinity than others. This study provided the predictions that these compounds may have the potential to prevent SARS-CoV-2 spike protein from entry into cells.

## Introduction

Recently, World Health Organization (WHO, https://www.who.int/) declared that the outbreak spread of the novel coronavirus severe acute respiratory syndrome coronavirus-2 (SARS-CoV-2) causes over 8 million reported cases and more than 4 hundred thousand deaths in 216 countries. Most of the coronaviruses cause only mild respiratory distress[1, 2]. However, three coronaviruses namely the severe acute respiratory syndrome coronavirus (SARS-CoV), SARS-CoV-2, and the Middle East respiratory syndrome coronavirus (MERS-CoV) have been found so far highly pathogenic transmitted from animal to human[3]. The SARS-CoV in 2003 brings about a 10 % case fatality rate (CFR), and MERS presented a CFR of about 34.4%[4-6]. The CFR for the novel coronavirus cannot be stated at the moment, as the pandemic is not over yet, although 3.4% is estimated and one thing is apparent the transmission rate of the infection is higher than the previous outbreaks[7].

The SARS-CoV-2 causes mainly severe acute respiratory distress by attacking lung cells and other complications in the heart, kidney, brain, and spleen and ultimately succumbed to a disastrous effect in coronavirus afflicted patients[8]. In this hour of crisis, there is no available effective standard-of-care present to treat the coronavirus affected patients. This makes an urgent need to find out agents to get control over the pandemic. Scientists globally make an effort to do their best to come up with effective therapeutic agents. So, to design a drug for the virus first we have to see the indispensable processes of the virus, which help them to survive and replicates its copy number. The genome of the virus showed a sequence similarity of 96.3% BatCoV RaTG13 and 79% with the SARS-CoV[9, 10]. SARS-CoV-2 is an enveloped single-stranded positive-sense RNA genome containing viruses. The RNA consists of 29,891 nucleotides, which transcribed for 6-12 ORFs, can be translated into around 28 identified non-structural and structural proteins (NCBI Reference Sequence: NC_045512.2)[11].

SARS-CoV-2 enters into the host cells containing ACE2, such as lung, oral, and nasal mucosa by glycosylated spike protein[12]. Human ACE2 interacts with SARS-CoV-2 spike proteins and facilitates SARS-CoV-2 entry into target cells[13]. The viral glycosylated spike protein composed of S1 and S2 subunits in the coronaviruses which enables them access into the host cell. The RBD of S1 subunit in SARS-CoV-2 attaches to ACE2, led to the shed of S1 subunit and subsequently triggers the cleavage of the S2 subunit by host protease protein, TMPRSS2, that can cleave S1/S2 protease cleavage site[13]. The cleavage changes the conformation and allowing HR1, HR2 to form 6-HB, which facilitates membrane fusion and virus releases its payload RNA into the cytoplasm. The first translation product of the RNA genome is polyprotein pp1a and pp1ab. Subsequently, 3CL protease (3CLpro) and papain-like protease (PLpro) cleave pp1a and pp1ab to functional proteins required for genome amplification. The other structural proteins are produced and the host ER-Golgi system assembles and releases mature virus from infected cells[14, 15]. The current understanding strongly recommends targeting the surface protein of the viral particle, which possibly prevents them from entering the host cell.

SARS-CoV-2 spike protein is a trimeric protein. The RBD of each subunit has two types of conformations. One conformation is “up” and the other is “down”. When the conformation is “up”, the viral spike protein could smoothly interact with human ACE2 protein, otherwise, it is inaccessible[16]. Comparing the structure of SARS-CoV-2 and SARS-CoV in spike protein, they have similar amino acid sequences and functions[17]. However, the RBD of SARS-CoV-2 spike protein in the down conformation N packs tightly against the N terminal domain (NTD) of the neighboring protomer, whereas the SARS-CoV-2 in the “down” conformation is angled closer to the central cavity of the trimer. Additionally, the binding affinity between SARS-CoV-2 spike protein and ACE2 protein is stronger than SARS-CoV, even 10-20 folds[18]. Therefore, we deduce that the structure of SARS-CoV-2 is similar to SARS-CoV, but not the same.

Drug discovery and development is a time taking, costly, and complex process. It usually takes years of effort to get clinically successful[19]. However, the virtual screening of authentic databases and the use of advanced bioinformatics and cheminformatics can reduce the time to come up with the best drug match to the selected target. This process of virtual screening has become a gold standard method for the preliminary phase of drug designing. To discover the potential hits against SARS-CoV-2, we have screened the essential entry pathway targeting virus penetration into cells. This study aims to find the potential hit compounds from Taiwan Database of Extracts and Compounds (TDEC, https://tdec.kmu.edu.tw/) for and against SARS-CoV-2 spike protein. We focus on the screen of hit compounds that can interact with viral spike protein and spike-ACE2 complex protein, leading to the prevention of viral payload into the host cytoplasm. We have selected our target mainly in the RBD of spike proteins for viral and host cell interactions. TDEC is an academic and scientific platform for investigators in different fields to share their research information. It includes much information, such as compounds’ structures, physicochemical properties, and biological activities, etc. on pure natural isolates, crude extracts, and synthesized extract from plants, microbes, marine organisms, and Chinese Herbal Medicines. In the present study, an attempt has been made to virtually screen candidates from TDEC for a selected target and did all the high throughput bioinformatics and cheminformatics analysis to get an insight into the interaction of spike protein and hits.

## Materials and methods

### Protein superimposition

The protein superimposition was performed by PyMOL software (version 0.99rc6) to compare the conformation of spike proteins in two states[20]. One was spike protein in the prefusion ACE2-free conformation, and the other was RBD of spike protein in the ACE2-bound conformation. In this study, the simulated structure of the RBD of spike protein in the prefusion ACE2-free conformation was constructed by homology modeling using the structure of the experimental structure of ACE2-free spike protein resolved by cryogenic electron microscopy (Cryo-EM) as the modeling template (S1A Fig) and the simulated spike protein was able to be obtained from SWISS MODEL website (S1B Fig)[18, 21]. The other was RBD of the spike protein bound with ACE2 protein that was an experimental protein structure resolved by X-ray crystallography and it was able to be downloaded from Protein Data Bank (PDB, https://www.rcsb.org/) (S1C Fig)[22]. The protein superimposition calculations were analyzed and shown by PyMOL software.

### Structure-based virtual screening

The virtual screenings with molecular docking calculations were performed by AutoDock Vina (version 1.1.2) program within PyRx (version 0.8) software to discover the potential compounds binding into the viral spike protein from the compound database [23, 24]. The 2,321 structures of compounds were downloaded (May 9, 2020) from TDEC, and then collected those compounds together and save it as an sdf format file by Discovery Studio 2019 visualizer software (DS 2019)[25]. Subsequently, these structures of compounds were optimized by energy minimization with steepest descent algorithm using MMFF94 force field, and then translated and divided to 2,321 individual pdbqt format files by Open Babel program within PyRx software[26]. The experimental structure of spike-ACE2 complex protein resolved by X-ray crystallography was obtained from Protein Data Bank (PDB ID: 6M0J) and the simulated structure of prefusional spike protein was downloaded from SWISS MODEL website[21, 22, 27]. After that, the substrates, including ligands, metal ions, and water molecules existed in both protein structures were all removed and the atoms of residues in both protein structures were modified by added the polar hydrogens and partial charges using DS 2019 software with CHARM force field. Moreover, to discover the potential compounds binding into the RBD of spike protein in the prefusion conformation and the site of the connective interface of spike-ACE2 complex protein from the database by virtual screening, dimensions of docking search spaces were respectively set big enough to contain the residues of ACE2 binding site on the RBD of spike proteins. Therefore, the size of the docking sites was respectively set as follows. In experimental spike-ACE2 complex protein resolved by X-ray crystallography, the coordinates of docking search space were respectively set at x = −38.6172, y = 28.4230, and z = 5.0328, and the size of dimensions of x, y, and z-axis (angstrom) were respectively set as x = 30.2373, y = 57.9713, and z = 19.4614. In other words, the docking space was set to the location which was at the connective interface of the spike-ACE2 complex protein. Besides, in the simulated spike protein in the prefusion conformation, the coordinates of docking search space were respectively set at x = 226.271, y = 195.386, and z = 306.4952, and the sizes of dimensions of x, y, and z-axis (angstrom) were respectively set as x = 47.2191, y = 38.0448, and z = 30.3288. It meant that the docking space was set to locate at the site on the prefusion RBD of the spike protein. Furthermore, the values of exhaustiveness used to search for molecular conformations were all set as 4 in docking calculation. Finally, the results of the docking simulations were shown and analyzed by PyMOL and DS 2019 software.

## Results

### Protein superimposition

A previous report had been shown that once the RBD of SARS-CoV-2 spike protein bound to human ACE2 protein, it can assist virus entry into the cells[17]. Recently, the 3D protein structure of the viral spike protein has been resolved by Cryo-EM technique[18]. However, we found that the structure of the spike protein resolved by Cryo-EM was incomplete because it lacked parts of residues in the RBD of spike protein (S1A Fig). Therefore, the structure of the spike protein with the complete amino acid sequences in the prefusion conformation was needed to be constructed by simulation. Fortunately, a useful and trusted simulated structure of SARS-CoV-2 spike protein in the prefusion conformation that has been constructed by homology modeling and could be easily downloaded from SWISS-MODEL website (S1B Fig). Additionally, the viral RBD of spike protein bound with human ACE2 protein, the spike-ACE2 complex protein, was also recently resolved by X-ray crystallography (S1C Fig). To investigate whether the conformation between the RBD of spike proteins in ACE2-free state and ACE2-bound state have differences or not, the protein superimposition was performed by PyMOL software using the structural aligning method. The data showed that the value of root mean square (RMS) of the two conformations was 1.735 Å (< 2Å). It meant that the superimposition between both RBD of spike proteins was good (Fig 1). However, comparing the RBD regions of both spike proteins in the simulation models, it was significantly different in structural conformation (blue ring in Fig 1). The simulation showed that the position of receptor-binding motif (RBM) regional structure in prefusion conformation was close to the RBD core than in the fusion conformation. Besides, the pose of RBM regional structure in the fusional conformation was more close to ACE2 protein than in the prefusion conformation. Generally, SAR-CoV-2 RBM contains many contacting residues binding to human ACE2 protein[22]. We concluded that the conformation changing of the RBM regional structure in the spike protein might affect the spike protein binding behavior to human ACE2 protein.

**Fig 1.**
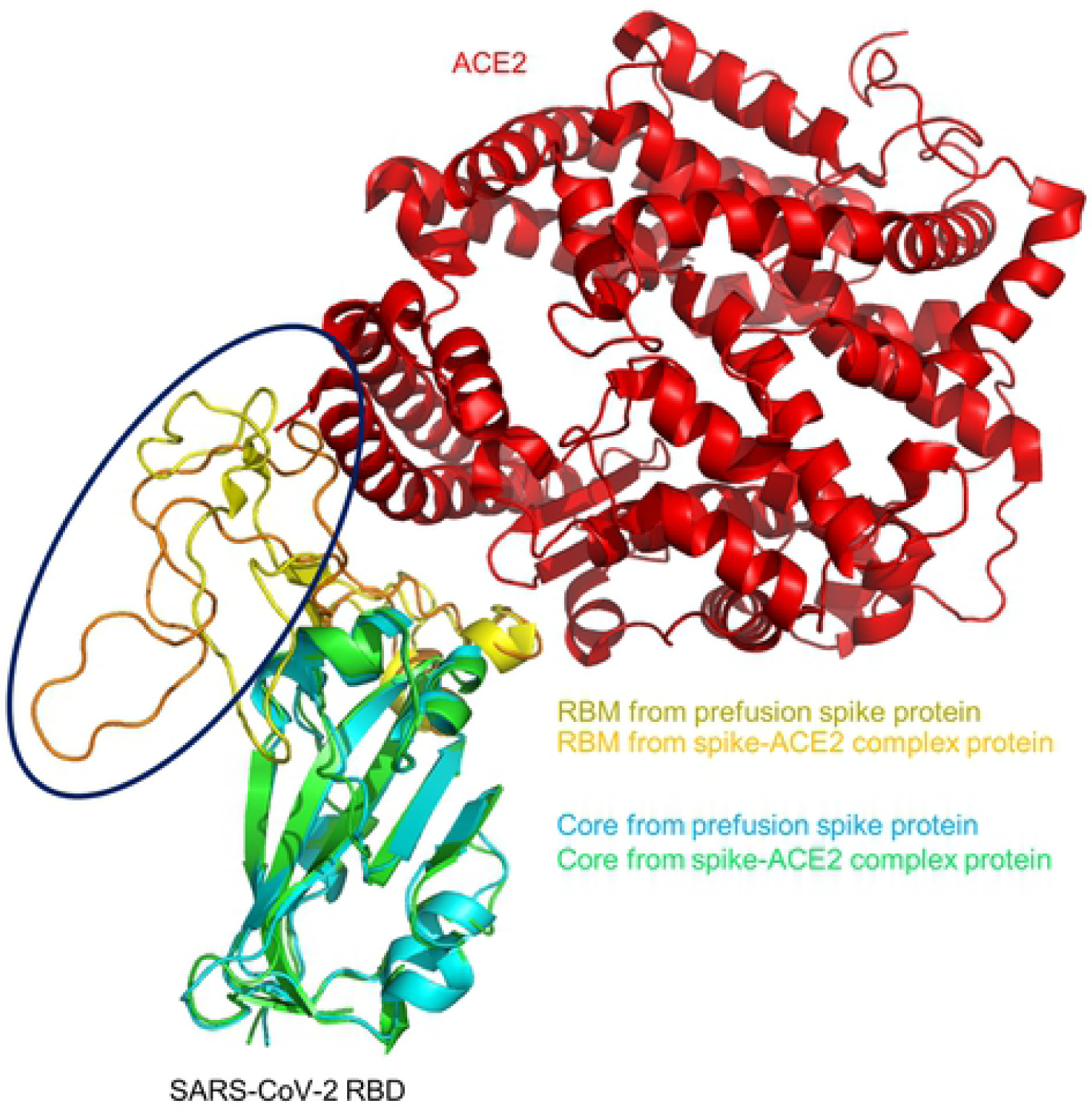
The protein superimposition of RBD of SARS-CoV-2 spike protein in two states. One is the simulated prefusional RBD protein constructed by homology modeling and the other is the experimental RBD protein of the spike-ACE2 complex protein resolved by X-ray crystallography. The SARS-CoV-2 RBD cores of the prefusional RBD structure and fusional RBD structure of the spike-ACE2 complex protein are respectively shown in cyan and green, and their RBM are respectively shown in orange and yellow. The ACE2 receptor is also shown in red. The major difference between the prefusional RBD of spike protein and the RBD of spike-ACE2 complex protein is shown within the blue ring.

### Structure-based virtual screening

The study reported that C-terminal domain (CTD) of SARS-CoV spike protein bound to ACE2 protein with different kinds of angles and its conformations or poses would be changed to well fit the conformation of dynamic ACE2 protein, such as ACE2 unbound-up conformation, ACE2-bound conformation, and ACE2 unbound-down conformation[10]. Additionally, we have observed that the conformation of the RBD of SARS-CoV-2 spike protein in the prefusional state is different from in the fusional state (Fig 1). In other words, we can speculate that the conformation of the dynamic RBD of SARS-CoV-2 spike protein would change to well bind and fit dynamic ACE2 protein. Therefore, we could design a strategy that two potential hitting sites could be set as the compound docking sites to prevent the spike protein to bind to ACE2 protein. One was that compounds docked on the RBD of the spike protein to prevent spike protein to bind to the human ACE2 protein, and the other was that compounds docked to the site of the connective interface of the spike-ACE2 complex protein to influence the affinity of binding between dynamic viral spike protein and dynamic human ACE2 protein and to cause nonfunctional changing of structural conformation to prevent viral spike protein from cleaved by human proteases. In this study, our aim was going to discover the compounds that not only can bind to the RBD of spike protein in the prefusion conformation but also bind to the site of the connective interface of spike-ACE2 complex protein. To discover the potential compounds, the virtual screening with molecular docking calculation was performed by AutoDock Vina program within PyRX software. All molecular structures prepared for virtual screening were downloaded from TDEC website. The results showed that the numbers of docked compounds (binding energy < −8 kcal/mol) in the prefusion RBD of spike protein and the spike-ACE2 complex protein were respectively 53 and 222. Among those compounds, the numbers of overlapped compounds and non-overlapped compounds were respectively 39 and 197. The binding energies and properties of 39 overlapped compounds were respectively shown in Fig 2, S2 Fig, and S1 Table. Subsequently, once the value of the screening threshold raised and limited to −9 kcal/mol, only compounds TDEC2018CN001781 (Thioflexibilolide A) and TDEC2020CN000246 (Candidine) reached the condition. Fig 3 and Table 1 respectively showed the structures and properties of the 2 compounds. Fig 4 showed the flowchart of virtual screening and provided a summary in this study.

**Table 1.** The list of properties of compounds TDEC2018CN001781 and TDEC2020CN000246.

| Property | Compound |  |
| --- | --- | --- |
| TDEC ID* | TDEC2018CN001781 | TDEC2020CN000246 |
| Name* | Thioflexibilolide A | Candidine |
| Source of Species* | *Sinularia flexibilis* | *Phaius mishmensis* |
| Molar mass [g/mol] * | 702.99 | 363.376 |
| Log P* | 6.54 | 3.88 |
| Solubility category* | Low | Low |
| Key of Interactive residues: |  |  |
| Pre-fusional Spike | Ala348, Leu452, Pro463, | Ile468, Leu492, |
|  | Leu492, Tyr451, Phe486 | Leu452, Phe486 |
| Spike-ACE2 Complex | Lys26, Asp30, Asn33,<br>His34 | Glu37, Asn33 |
| Binding Energy (kcal/mol): |  |  |
| Prefusion Spike | -9.2 | -9.0 |
| Spike-ACE2 Complex | -8.9 | -9.8 |

\*The data were obtained from TDEC

**Fig 2.**
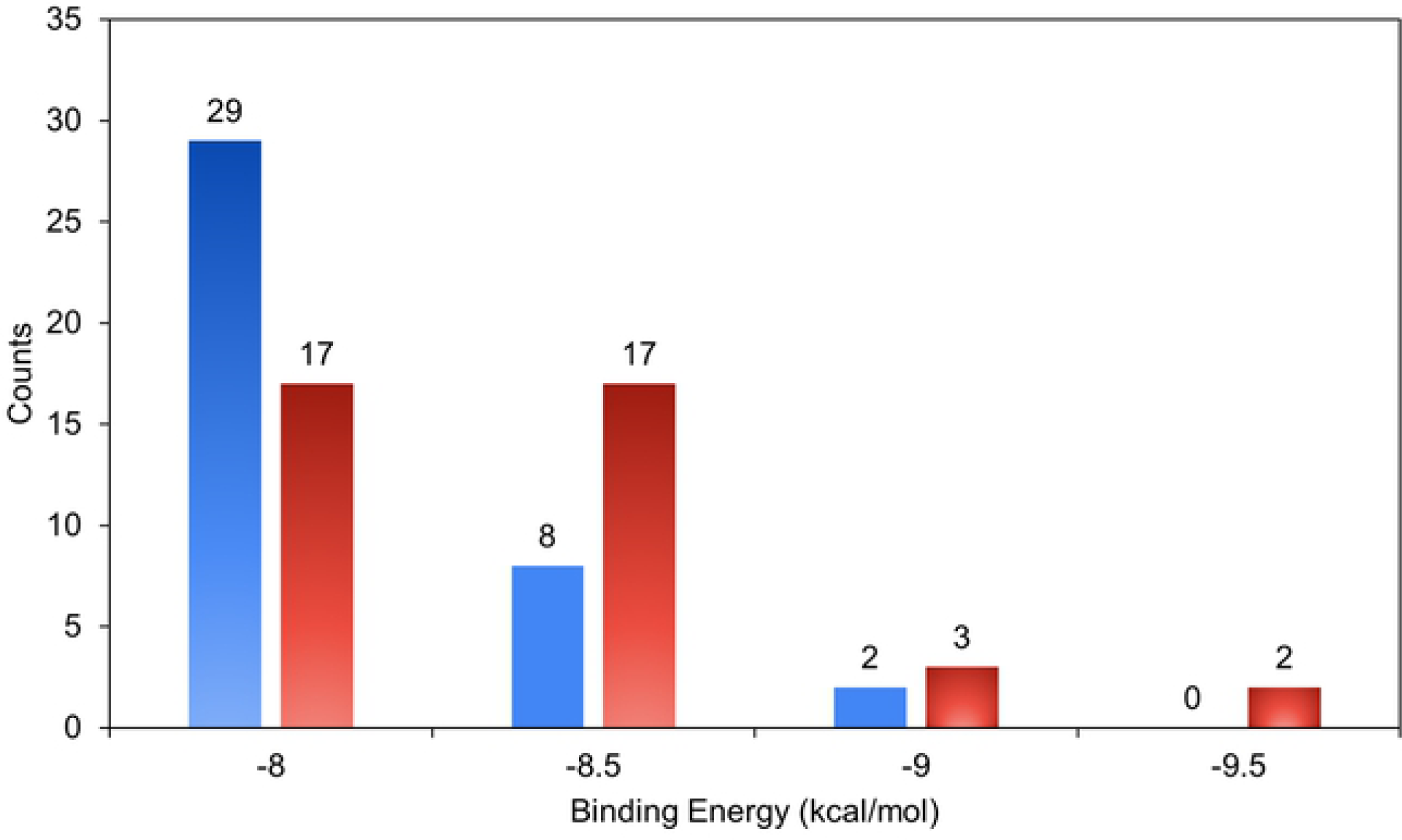
The values of binding affinities of 39 overlapped potential compounds. The blue bar showed the numbers of compounds docking to the prefusional spike protein and the other red bar showed the numbers of compounds docking to the ACE2-spike complex protein. Note, all values of binding energy in the 39 potential compounds were less than −8 kcal/mol.

**Fig 3.**
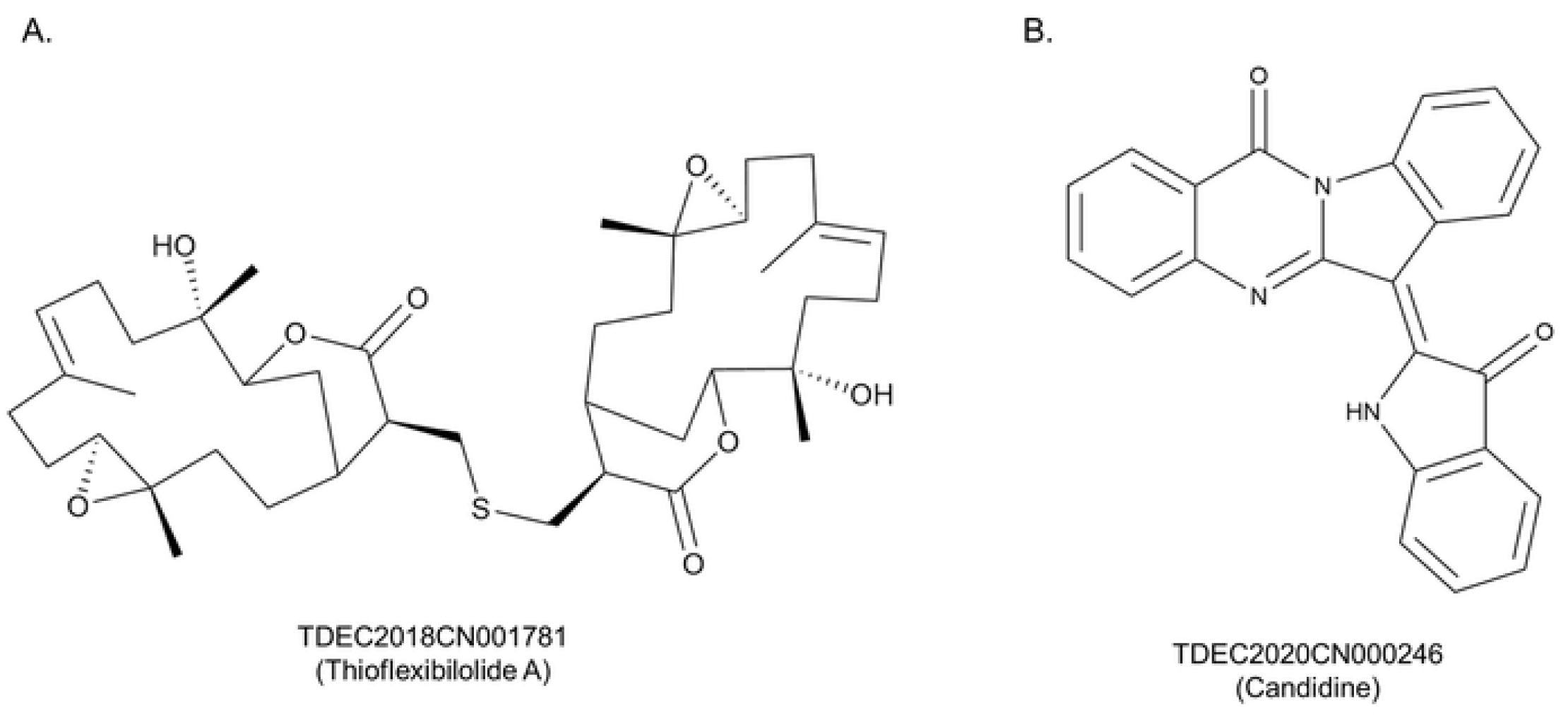
The structures of Thioflexibilolide A and Candidine. (A) TDEC2018CN001781 (Thioflexibilolide A) and (B) TDEC2020CN000246 (Candidine)

**Fig 4.**
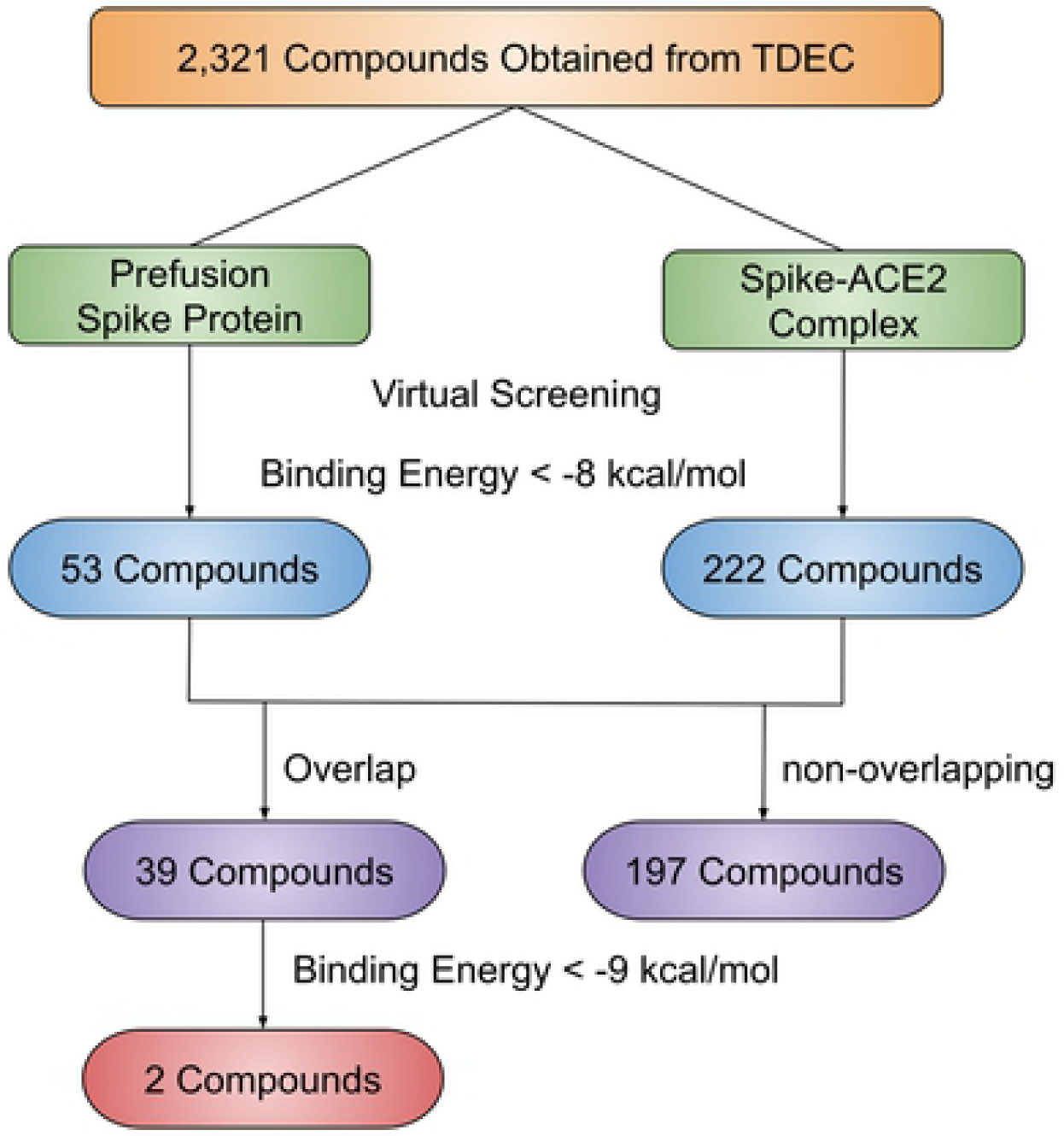
The flowchart of the structure-based virtual screening.

The data showed that thioflexibilolide A (yellow stick) could dock with the RBD of spike protein (cyan cartoon) and its value of binding energy was −9.2 kcal/mol (Fig 5A). It binds to the target site and forms many interactions with amino acids in the prefusional RBD of spike protein. The results showed that it formed six alkyl-alkyl interactions (purple dashed line) with amino acids Ala348, Leu452, Pro463, and Leu492, and their interactive distances were respectively 4.43Å, 4.85Å, 3.63Å, 4.00Å, 4.96Å, and 5.12Å. In addition to that, it also formed two pi-alkyl interactions (light purple dashed line) with amino acids Tyr451 and Phe486, and their interactive distances were respectively 4.29Å and 4.52Å (Fig 5B). Moreover, thioflexibilolide A also successfully docked with the site near the connective interface of the RBD of spike-ACE2 complex protein and its value of binding energy was −8.9 kcal/mol (Fig 5C). Besides, it also formed many interactions with amino acids in the space near the connective interface of the RBD of spike-ACE2 complex protein. The results showed that thioflexibilolide A formed three hydrogen-bonding interactions (green dashed line) with amino acids Lys26, Asp30, and Asn33, and their interactive distances were respectively 2.11Å, 2.87Å, and 1.81Å and an alkyl-alkyl interaction (purple dashed line) with amino acid Lys26, and its interactive distance was 2.87Å. Moreover, it also formed a pi-alkyl interaction (light purple dashed line) with amino acid His34, and its interactive distance was 4.71Å (Fig 5D).

**Fig 5.**
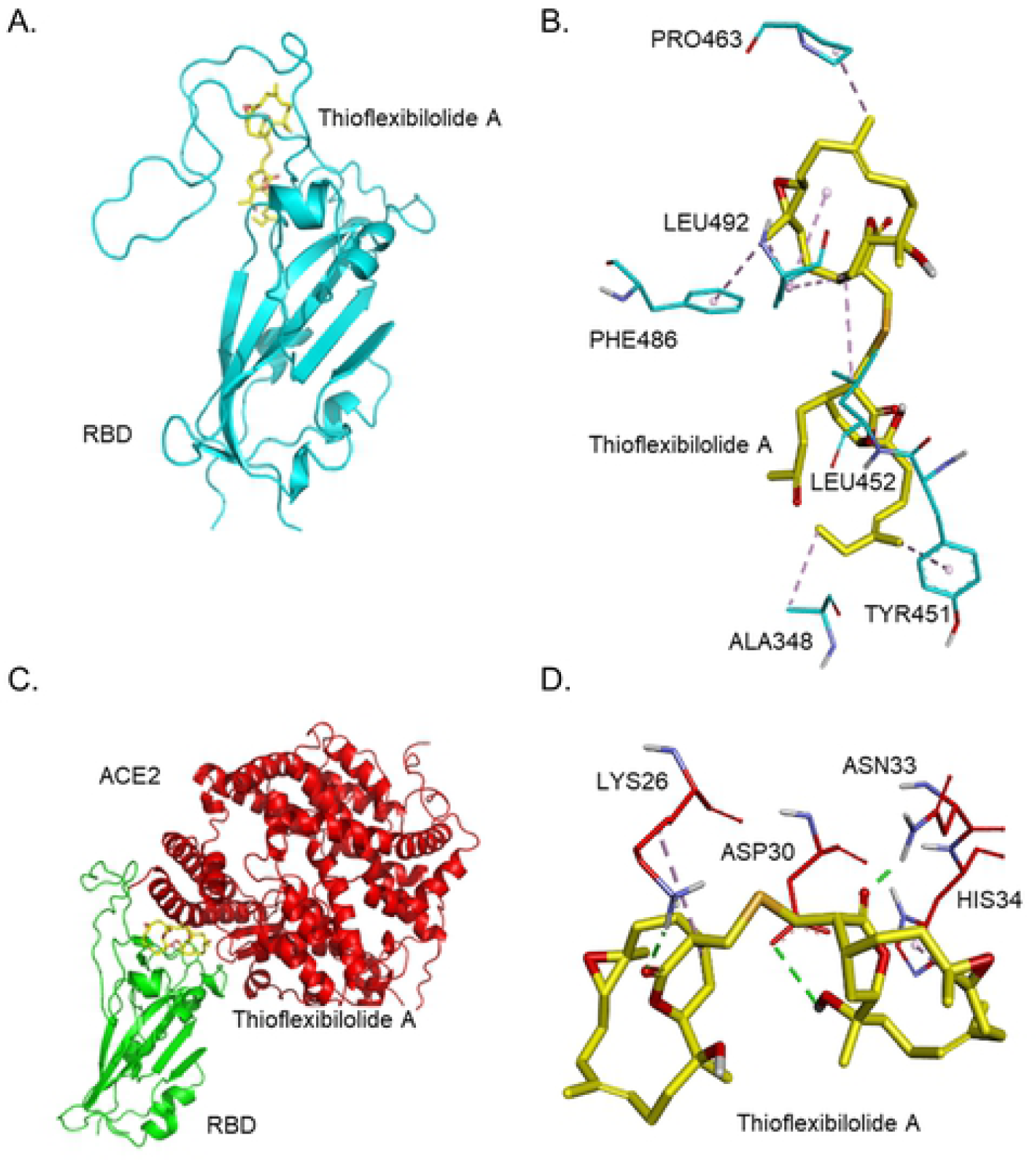
The simulation of thioflexibilolide A docking with SARS-CoV-2 spike protein. (A) Thioflexibilolide A (yellow stick) docked with the prefusional RBD of spike protein (cyan cartoon). (B) Thioflexibilolide A formed eight hydrophobic interactions (purple dashed line) with amino acids (cyan stick) in the RBD of the spike protein. (C) Thioflexibilolide A docked with the site near the connective interface of the spike-ACE2 complex. The RBD of spike protein and ACE2 respectively shown in the green and red cartoons. (D) Thioflexibilolide A formed two hydrophobic interactions (purple dashed line) and three electrostatic hydrogen-bonding interactions (green dashed line) with amino acids (red stick) near the connective interface of spike-ACE2 complex protein.

The data also showed that candidine could dock with the prefusional RBD of spike protein (cyan cartoon) and its value of binding energy was −9.0 kcal/mol (Fig 6A). It also formed many interactions with amino acids in the prefusional RBD of the spike protein. The results showed that it formed two pi-alkyl interactions (light purple dashed line) with amino acids Ile468 and Leu492, and their interactive distances were respectively 4.86Å and 4.67Å and four pi-sigma interactions (dark purple dashed line) with amino acid Leu452, and their interactive distances were respectively 3.83Å, 3.95Å, 3.77Å, and 3.55Å. Moreover, it also formed a pi-pi T-shaped interaction (pinking purple dashed line) with amino acid Phe486, and its interactive distance was 5.39Å (Fig 6B). Candidine docked with the site near the connective interface of the RBD of spike-ACE2 complex protein and its value of binding energy was −9.8 kcal/mol (Fig 6C). Besides, it also formed many interactions with amino acids in the space near the connective interface of the RBD of spike-ACE2 complex protein. The results showed that it formed an electrostatic hydrogen-bonding interaction (green dashed line) with amino acid Glu37, and its interactive distance was 2.25Å. Besides, it also formed a hydrophobic pi-sigma interaction (dark purple dashed line) with amino acid Asn33, and its interactive distance was 3.94Å (Fig 6D).

**Fig 6.**
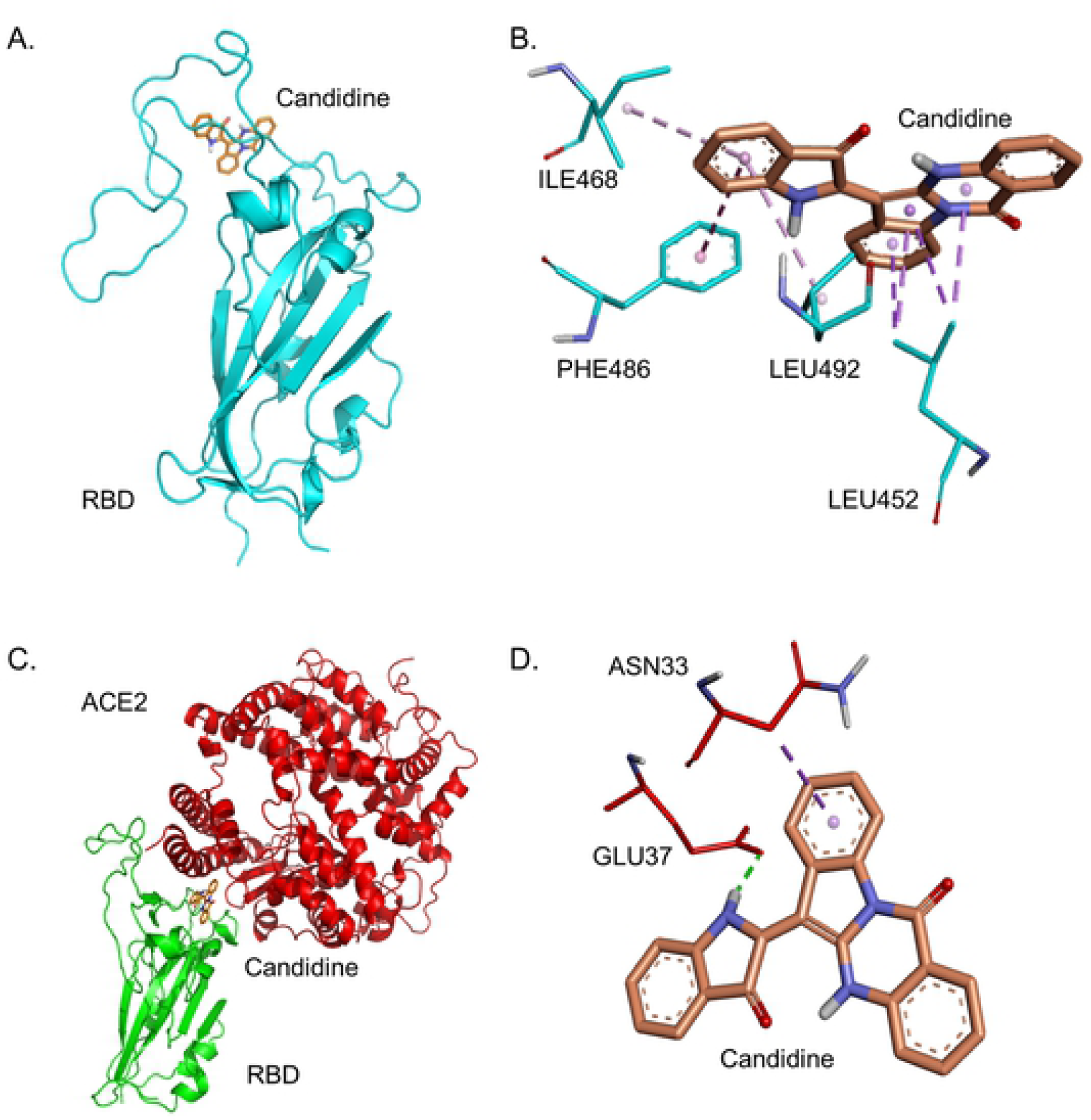
The simulation of candidine docking with SARS-CoV-2 spike protein. (A) Candidine (orange stick) docked with the prefusional RBD of spike protein (cyan cartoon). (B) Candidine formed seven hydrophobic interactions (purple dashed line) with amino acids (cyan stick) in the RBD of the spike protein. (C) Candidine docked with the site near the connective interface of spike-ACE2 complex. The RBD of spike protein and ACE2 respectively shown in the green and red cartoons. (D) Candidine formed an electrostatic interaction (green dashed line) and hydrophobic interaction (purple dashed line) with amino acids near the connective interface of spike-ACE2 complex protein.

In conclusion, the results revealed that two natural products thioflexibilolide A and candidine may have the potential abilities to bind to the RBD of spike protein in the prefusion conformation and spike-ACE2 complex protein and have good binding affinities than others. In other words, the compounds may have the potential efficacy to prevent the viral spike protein from binding to human ACE2 protein. Moreover, some residues located in the two proteins might play key roles in the compound binding. The simulation showed that both compounds could interact with the same amino acids Leu452, Phe486, and Leu 492 in the prefusion conformation of RBD of spike protein, and the same amino acid Asn33 in the spike-ACE2 complex protein.

## Discussion

COVID-19 becomes difficult now to manage COVID-19 patients because of the absence of an effective vaccine as well as drug availability. Hence, there is an expedite need for new drug development against SARS-CoV-2. Previous studies have been reported many virtually screening drugs targeting proteases, such as 3CL protease (3CLpro) and papain-like protease (PLpro), which processed the essential polyprotein of infectious coronavirus[17, 28, 29]. The present study aimed to first screen the hit compounds for SARS-CoV-2 spike protein. We have aimed the conformations of target sites from viral surface protein, the prefusional spike protein and fusional spike-ACE2 complex protein for our virtual screening method to identify the best possible drugs against SARS-CoV-2. The target sites are vital at the entry point of viral particles entering into the cell of the host. As the functional spike protein and its interaction with ACE2 is substantial because it governs virus entry of the virus into the host cell, this makes it an attractive target in developing drugs and vaccines for coronaviruses[13, 30]. The predictions obtained from virtual screening such as binding energy, possible flexibility in conformation, are still very useful to lead the way of drug discovery or drug repurposing. As represented in Fig 4, we have started with 2,321 compounds taken from TDEC and applied the filters to get compound which can bind to the target structures with high affinity. Notably, we got the two natural products, thioflexibilolide A and candidine, which passed all the filters applied with the strong binding energy respectively −9.2 kcal/mol and −9.0 kcal/mol on the prefusion spike protein and −9.8 kcal/mol and −8.9 kcal/mol on the spike-ACE2 complex protein (Fig 5 and Fig 6). The other compounds listed in the S1 Table have reasonable binding energy around −8 kcal/mol, which can be considered in the further phenotypic assay. Among them, hit compounds were able to influence the interaction between viral RBD of the spike protein and the human ACE2 protein in two dynamic proteins. Besides, we also found among these 39 compounds that Dioscin, Actinomycin D and Saikosaponin C had been reported that they had anti-virus activity against others virus (S2 Table). Although, those compounds that we discovered from the database may have the potential ability to prevent the binding between viral spike protein and human ACE2 protein, the efficacy of compounds were still needed to be further verified by bioassay.

Thioflexibilolide A and candidine were natural products respectively isolated from soft coral *Sinularia flexibilis* and *Phaius mishmensis*[31, 32]. In chemical structure, the structure of candidine is more solid than thioflexibilolide A. It suggested that candidine could be more specificity than thioflexibilolide A in binding ability with viral spike protein. Additionally, the docking model also showed that the binding positions of thioflexibilolide A and candidine were similar whatever in the prefusional RBD of spike protein or spike-ACE2 complex protein. The results showed that both compounds could interact with the same residues whatever located in the prefusional RBD of spike protein or the connective interface of spike-ACE2 complex protein. Furthermore, in the docking models, the results also showed that both compounds bound to the side face of RBM of the spike protein. We deduced that it might be caused that the RBM of spike protein is major composed of chain type of structure rather than α-helix or β-sheet structure (S1 Fig). Besides, we also deduce that two compounds bind to side face respectively with the RBM of the ACE2-free spike protein and the connective interface of spike-ACE2 complex protein probably affect the affinity of binding between viral spike protein and human ACE2 protein or influence its conformation changing to prevent spike protein to be cleaved by host protease like TMPRSS2 (Fig 5 and Fig 6).

To rapidly develop useful drugs against SARS-CoV-2, computer-aid drug screening was performed in many studies. For instance, Chen *et al*. reported that drugs Epclusa (velpatasvir/sofosbuvir) and Harvoni (ledipasvir/sofosbuvir) may have the inhibition for viral 3CLpro**[13]** by virtual screening. Besides, Ma *et al*. screened 12,322 and 11,294 potential components respectively against viral 3CLpro and PLpro from TCMD 2009 database *in silico*[33]. Furthermore, Khana *et al*. respectively identified 2 potential compounds for viral 3CLpro and 20-O-ribose methyltransferase from an in-house library of 123 antiviral drugs by virtual screening[28]. Moreover, Ton *et al*. found 1,000 potential compounds for viral 3CLpro that may have the efficacy against SARS-CoV-2 from ZINC15 library by structure-based virtual screening using Deep Docking[34]. Additionally, Gentile discovered 17 potential inhibitors against viral 3CLpro from marine natural products by virtual screening integrating with pharmacophore and molecular dynamic[30]. Besides, Wang identified drugs Carfilzomib, Eravacycline, Valrubicin, Lopinavir, and Elbasvir had good binding affinities for 3CLpro by virtual screening integrating with docking calculation and molecular dynamics simulations[35]. Likewise, Pant *et al*. utilized *in-silico* approaches screened 300 potential compounds against viral 3CLpro from CHEMBL database, ZINC database, FDA approved drugs database[36]. Liu *ex al*. found an anticoagulation agent dipyridamole (DIP) from U.S. Food and Drug Administration (FDA) approved drug library by virtual screening. The literature also pointed out that it could suppress SARS-CoV-2 replication *in vitro*[37]. Sarma *et al*. identified 2 compounds against viral NTD of nucleocapsid protein (N protein) from Asinex and Maybridge library by virtual screening integrating with docking, molecular property, and molecular dynamic[29].

Wu *et al*. reported for the first time that screening potential compounds for SARS-CoV-2 spike protein from compound databases by virtual screening [38]. They constructed 20 homology structures including 19 viral targets and 1 human target and screened many potential compounds from ZINC database, in-house natural products, and 78 antiviral drugs by virtual screening and fast shared their huge results to researchers around the world in February of this year. At that time, structures of both SARS-CoV-2 spike protein and spike-ACE2 complex protein yet be resolved and published. Therefore, based on the reasons, they constructed the simulated RBD of SARS-CoV-2 spike protein by homology modeling using SARS-CoV spike glycoprotein (PDB ID: 3SCI) as the template and built spike-ACE2 complex protein by protein-protein docking. Recently reported had pointed out that SARS-CoV-2 and SARS-CoV spike protein were homology proteins with greater than 70% similar sequences and have the same function for binding ACE2. However, the previous report also showed that the binding affinity between SARS-CoV-2 spike protein and ACE2 was higher than SARS-CoV spike protein approximately 10-20 folds[18, 39]. These reports suggested that it is highly similar but not the same in partial structure or conformation in the spike proteins between SARS-CoV-2 and SARS-CoV. Moreover, spike-ACE2 complex protein was constructed by protein-protein docking may not good enough to directly exhibit the protein-protein binding situation in fact. Recently, the RBD of SARS-CoV-2 spike protein in the prefusion conformation (PDB ID: 6VSB) and the complex protein that SARS-CoV-2 spike protein’s RBD bound with ACE2 protein had already been resolved (PDB ID: 6M0J) respectively by Cryo-EM and X-ray crystallography with high resolution[18, 22]. Therefore, the two new resolving protein structures were likely more close to the structural situations in fact. In other words, it could raise the accuracy in structure-based virtual screening with docking calculation because of those structures resolved with high resolution that is why we did that in this study.

Developing the viral spike antibody is one of the good strategies to fight against SARS-CoV-2. It could bind to the RBD of spike protein to neutralize SARS-CoV-2 had been reported [40, 41]. However, it is concerned about inducing antibody-dependent enhancement (ADE)[42]. Herein, we reported that the 39 potential compounds against SARS-CoV-2 spike protein may have the advantage of avoiding to induce ADE.

ACE2 was a type I transmembrane protein and it functioned as a carboxypeptidase to mediate homeostasis of renin-angiotensin system (RAS), such as blood volume, lung and cardiovascular regulating functions, by cleaving angiotensin II (Ang II) to Ang(1-7)[43]. ACE2 was higher expressed in the small intestine, kidneys, testicles, adipose tissue, and thyroid and moderate in the lungs, liver, adrenal glands, and rectum and lower in the blood, blood vessels, spleen, brain, muscle, and bone marrow[44, 45]. On the other hand, gender and age characters could affect ACE2 expression under inflammatory conditions in the skin, lung, brain, and thyroid[44]. Recent studies also showed that gender could affect SARS-CoV-2 infection susceptibility and virus clearance[17]. Besides, another report also pointed out that COVID-19 patients’ symptoms including dysfunctions in diarrhea, taste, and smell; injuries in the liver, kidney, and heart may have a close relationship with ACE2 expression[46]. Herein, our results showed the 39 potential compounds could prevent the binding between viral spike protein and human ACE2 protein *in silico*. It is speculated that those potential compounds could fight against COVID-19 and lower the expression in the above symptoms for humans.

Natural products proved to be beneficial due to the contribution from a long time to develop effective drugs for several diseases[47]. Therefore, we have tried our endemic TDEC which is very rich and diverse in natural product resources derived from traditional medicine, domestic microbes, and marine organisms. thioflexibilolide A is well-known for its neuroprotective and anti-inflammatory activities derived from soft corals, whereas about gravicycle is reported non-toxic after its *in vitro* toxicity assessment in the cell[48]. The results we are reporting here about the compounds which are screened by employing virtual screening with molecular docking approach from the database for disabling SARS-CoV-2 spike protein’s RBD from its interaction with ACE2 receptor of the host cell is very first of its kind as far as possibly know. The compounds perform well to get the filter into potential drug candidate during our screening process, and we strongly recommend our compound for further *in vitro* and *in vivo* investigation.

## Conclusion

The study carried out intending to come up with potential drugs for COVID-19 therapy. Here in this piece of work, we virtually screened more than 2000 drugs against the target spike protein’s RBD. After data mining and filtering out the unfitted compounds, we finally found two compounds named thioflexibilolide A and candidine. We have done the molecular docking to check whether it binds with the target with strong affinity or not. As per our result, we found these two compounds bind with strong affinity to the selected target of the viral protein. We believed that these two compounds could be potential drugs for SARS-CoV-2. The two compounds needed further cell-based validation and could be a hope to develop anti-SARS-CoV-2 therapy.

## Acknowledgments

We thank the Taiwan Database of Extracts and Compounds website for the assistance of offering chemical and biological information of extracts and compounds. Besides, we are grateful to the National Center for High Performance Computing for computer time and facilities. We also thank the Center for Resources, Research and Development of Kaohsiung Medical University for the ChemBio3D Ultra 11.0 technical support.

## Supporting information

**S1 Fig. The structures of SARS-CoV-2 spike proteins in different conditions**. (A) Spike protein composed of trimers (green, cyan, pink cartoons) in the prefusion conformation resolved by cryogenic electron microscopy (Cryo-EM) and could be downloaded from Protein Data Bank (PDB ID: 6VSB)[18]. (B) Structure of the spike protein simulated by homology modeling. The homology modeling spike protein structure using the viral spike protein structure which was resolved by Cryo-EM as the modeling template and could be obtained from SWISS MODEL website[18, 21]. The quality of simulated spike protein is good (GMQE score = 0.72 (0 < score < 1), GMEAN score = −2.81 (score > −4.0)) that verification was provided by SWISS MODEL[49, 50] (C) Structure of spike-ACE2 complex protein was resolved by X-ray crystallography and obtained from PDB (PDB ID: 6M0J)[22]. The RBD of spike protein and ACE2 respectively shown in the green and red cartoons. (D) Compared the RBD of spike proteins in the prefusion conformation between the structure which was resolved by Cryo-EM (yellow cartoon) and the simulated structure (green cartoon) by protein superimposes using PyMOL software. The data showed the value of root mean square (RMS) of the two conformations was 0.713 Å (< 2Å). On the other hand, the structural superimpose between both proteins was good. The structures in the region within the blue frame are the range of lacking residues and would be filled by homology modeling.

**S2 Fig. The structures of 39 potential compounds**. The 39 compounds (binding energy < −8) that not only can bind to the RBD of spike protein in the prefusion conformation but also bind to the site of the connective interface of spike-ACE2 complex protein. The TDEC ID of each structure was also shown in figure.

**S1 Table.** The list of 39 potential compounds. The list showed the information of each compound, including TDEC ID, binding energy in prefusion spike protein and spike-ACE2 complex, and species source.

| Compound | Binding energy (kcal/mol) |  | Species source |
| --- | --- | --- | --- |
|  | Prefusion spike protein | Spike-ACE2 complex protein |  |
| TDEC2018CN001781 | -9.2 | -8.9 | *Sinularia flexibilis*[31] |
| TDEC2020CN000246 | -9.0 | -9.8 | *Phaius mishmensis*[32, 51] |
| TDEC2019CA001707 | -8.9 | -8.3 | *Tripterygium wilfordii*[52] |
| TDEC2019CA001572 | -8.8 | -8.9 | *Dioscorea nipponica*[53] |
| TDEC2019CN000617 | -8.8 | -8.3 | *Solanum torvum*[54] |
| TDEC2018CN001906 | -8.7 | -8.0 | *Sinularia crassa*[55] |
| TDEC2018CN000709 | -8.6 | -8.1 | *Streptomyces sindenensis*[56] |
| TDEC2019CA001664 | -8.6 | -9.1 | *Bupleurum chinense*[57] |
| TDEC2019CA001644 | -8.5 | -9.1 | *Cunninghamia lanceolata*[58] |
| TDEC2019CN000295 | -8.5 | -9.3 | *Solanum torvum*[59] |
| TDEC2018CA000824 | -8.4 | -8.2 | *Strongylophora durissima*[60] |
| TDEC2018CN001905 | -8.4 | -8.4 | *Sinularia crassa*[55] |
| TDEC2018CN002006 | -8.4 | -9.5 | *Sinularia brassica*[61] |
| TDEC2019CN000357 | -8.4 | -8.9 | *Paraminabea* *acronocephala* [62] |
| TDEC2019CN000432 | -8.4 | -8.3 | *Sapindus mukorossi*[63] |
| TDEC2019CN000548 | -8.4 | -8.8 | *Calamus*<br>*quiquesetinervius*[31] |
| TDEC2018CA000891 | -8.3 | -8.6 | *Clerodendrum inerme*<br>derivatives[64] |
| TDEC2018CN001829 | -8.3 | -8.9 | *Paraminabea*<br>*acronocephala*[62] |
| TDEC2018CN002007 | -8.3 | -8.3 | *Sinularia brassica*[61] |
| TDEC2019CN000274 | -8.3 | -8.8 | *Solanum torvum*[65] |
| TDEC2020CN000050 | -8.3 | -8.1 | *Artocarpus communis*[66] |
| TDEC2018CN002164 | -8.2 | -8.0 | *Sinularia brassica*[64] |
| TDEC2019CA001733 | -8.2 | -8.1 | *Epimedium*<br>*brevicornum*[67] |
| TDEC2019CN000275 | -8.2 | -8.5 | *Solanum torvum*[68] |
| TDEC2019CN000476 | -8.2 | -8.8 | *Perrottetia arisanensis*[69] |
| TDEC2019CN000616 | -8.2 | -8.0 | *Lawsonia inermis*[70] |
| TDEC2019CN000634 | -8.2 | -8.1 | *Solanum violaceum*[71] |
| TDEC2019CN000635 | -8.2 | -8.4 | *Solanum violaceum*[71] |
| TDEC2018CN001495 | -8.1 | -8.2 | *Isis hippuris*[72] |
| TDEC2019CN000249 | -8.1 | -8.9 | *Sapindus mukorossi*[73] |
| TDEC2019CN000268 | -8.1 | -8.5 | *Solanum torvum*[68] |
| TDEC2019CN000674 | -8.1 | -8.1 | *Rutaceae* derivatives[74]<br>[75] |
| TDEC2020CN000221 | -8.1 | -8.7 | *Cephalantheropsis gracili*[76, 77] |
| TDEC2018CN002008 | -8.0 | -8.8 | *Sinularia brassica*[61] |
| TDEC2018CN002161 | -8.0 | -8.6 | *Sinularia brassica*[61] |
| TDEC2019CN000527 | -8.0 | -8.7 | *Annona montana*[63] |
| TDEC2019CN000665 | -8.0 | -8.5 | *Paraminabea acronocephala*[62] |
| TDEC2019CN000667 | -8.0 | -8.1 | *Paraminabea acronocephala*[62] |
| TDEC2019CN000203 | -8.0 | -8.6 | *Sapindus mukorossi*[63] |

**S2 Table.** The list showed the compounds which had been reported having anti-virus activity in the literatures. The information of logP and solubility category were all obtained from TDEC.

| Ligand | Name | logP | Solubility category | Application |
| --- | --- | --- | --- | --- |
| TDEC2019CA001572 | Dioscin | 1.71 | Moderate | Anti-virus[78, 79] |
| TDEC2018CN000709 | Actinomycin D | -0.1 | Moderate | Anti-virus[56, 80, 81] |
| TDEC2019CA001664 | Saikosaponin C | -0.37 | High | Anti-virus[82, 83] |

